# Transcranial photoacoustic imaging of NMDA-evoked focal circuit dynamics in rat hippocampus

**DOI:** 10.1101/308585

**Authors:** Jeeun Kang, Shilpa D. Kadam, Joshua S. Elmore, Brennan J. Sullivan, Heather Valentine, Adarsha P. Malla, Maged M. Harraz, Arman Rahmim, Jin U. Kang, Leslie M. Loew, Michael Baumann, Anthony A. Grace, Albert Gjedde, Emad M. Boctor, Dean F. Wong

## Abstract

Transcranial functional photoacoustic (fPA) voltage-sensitive dye (VSD) imaging promises to overcome current temporal and spatial limitations of current neuroimaging modalities. The technique previously distinguished global seizure activity from control neural activity in groups of rats. To validate the focal specificity of transcranial fPA neuroimaging *in vivo*, we now present proofs-of-concept that the results differentiate between low- and high-dose N-methyl-D-aspartate (NMDA) evoked neural activity in rat hippocampus. Concurrent quantitative EEG (qEEG) and microdialysis recorded real-time circuit dynamics and glutamate concentration change, respectively. We hypothesized that location-specific fPA VSD contrast would identify the neural dynamics in hippocampus with the correlation to NMDA evoked focal glutamate release and time-specific EEG signals. To test the hypothesis, we infused 0.3 to 3.0 mM NMDA at 2 μl/min over 60 min via an implanted microdialysis probe. The dialysate samples collected every 20 min during the infusion were analyzed for focal changes in extracellular glutamate release. Transcranial fPA VSD imaging provided NMDA-evoked VSD responses with positive correlation to extracellular glutamate concentration change at the contralateral side of the microdialysis probe. The graded response represents the all-or-none gating system of the dentate gyrus (DG) in hippocampus. Quantitative EEG (qEEG) successfully confirmed induction of focal seizure activity during NMDA infusion. We conclude that transcranial fPA VSD imaging distinguished graded DG gatekeeping functions, based on the VSD redistribution mechanism sensitive to electrophysiologic membrane potential. The results suggest the potential future use of this emerging technology in clinics and science as an innovative and significant functional neuroimaging modality.

## Introduction

Whereas electrophysiological and invasive neurochemical techniques have been valuable in assessing activity within specific neural circuits in brain, there is a distinct need for a non-invasive means to image real-time regional functional activity in deep brain *in vivo*. The hippocampal circuit has rows of excitatory pyramidal neurons all aligned in close proximity with similar orientation. The dentate gyrus (DG) is classically considered as a gate keeper for the propagation of excitatory activity into the hippocampus as the excitatory neurons of the DG are less excitable than other types of hippocampal neurons. Therefore, low amplitude excitatory inputs do not generate action potentials in DG neurons and fail to open the DG gate ^1^. In contrast to sub-threshold excitatory stimulus, strong repetitive excitatory inputs can break the DG gate and induce seizure activity ^1^. This all-or-none gating function of the DG is consistent with the hippocampus being the lowest seizure threshold region of the mammalian brain.

Previous optical imaging approaches to local cortical circuits ^2-4^ applied invasive procedures to overcome superficial imaging depth, and further extension to deeper brain regions such as the hippocampus was prohibitive. The imaging of electrophysiological or neurochemical dynamics within the hippocampus primarily has been accomplished by optical imaging of freshly-sliced brain tissue ^5^ or magnetic resonance imaging spectroscopy (MRS) for non-invasive quantification of glutamate at high spatial resolution ^6,7^. However, optical neuroimaging and MRS suffer from shallow imaging depth and from slow imaging speed, respectively. The application of two-photon microscopy to measure calcium ion dynamics in specific deep brain structures, including hippocampus, has poor temporal resolution ^8^. Therefore, non-invasive, real-time neuroimaging modality having a high spatiotemporal resolution would be a significant advance, but such an approach would require neurochemical and electrophysiological characterization to validate the neural activity.

Photoacoustic (PA) imaging is a hybrid approach combining optics and acoustics where the signal corresponding to the neural activity is detected in the form of acoustic transcranial imaging with optical absorbance as an image contrast ^9,10^. PA imaging is based on thermo-elastic perturbation of a target evoked by light absorbance from a pulsed laser illumination, which generates radio-frequency (RF) acoustic pressure waves detected by piezoelectric ultrasound transduction or optical interferometry. With this unique imaging mechanism, several attractive applications have been proposed for preclinical and clinical research with tomographic and microscopic imaging modes, including detection of endogenous contrast of cancer indicators, e.g., melanoma ^11^, breast microcalcifications ^12-14^, monitoring of cancer angiogenesis ^15^, oxygen metabolism ^16^, and quantification of lipid content ^17,18^, among others. Recently, we presented the functional PA (fPA) neuroimaging of near-infrared VSD redistribution mechanism differentiating the graded membrane potential in lipid vesicle model and chemo-convulsant seizure in rodent brain *in vivo* ^19^.

Here, we validated the transcranial fPA neuroimaging of hippocampal electrophysiology activated by stimulation of N-methyl-D-aspartate (NMDA) receptors in rat brain *in vivo* through intact skull and scalp. The tri-modal neural sensing approach utilized microdialysis, quantitative electroencephalography (qEEG), and transcranial fPA neuroimaging to simultaneously monitor electrophysiological changes associated with glutamatergic neurotransmission (Figure 1). To stimulate the hippocampus, we infused NMDA by reverse microdialysis that triggered localized changes of extracellular glutamate concentration. We quantified the change of extracellular glutamate concentration evoked by reverse microdialysis by direct microdialysis and high-performance liquid chromatography (HPLC). We combined fPA neuroimaging and qEEG with microdialysis to evaluate the effects associated with graded DG gatekeeping at hippocampus, in the presence of glutamatergic excitation. Concurrently, the electrophysiological signatures were captured with transcranial fPA neuroimaging. Each of the three modalities thus provided information with different degrees of specificity, including spatiotemporal specificity from fPA imaging, temporal specificity from qEEG, and neurochemical specificity from microdialysis in the target neural circuit.

**Figure 1.**
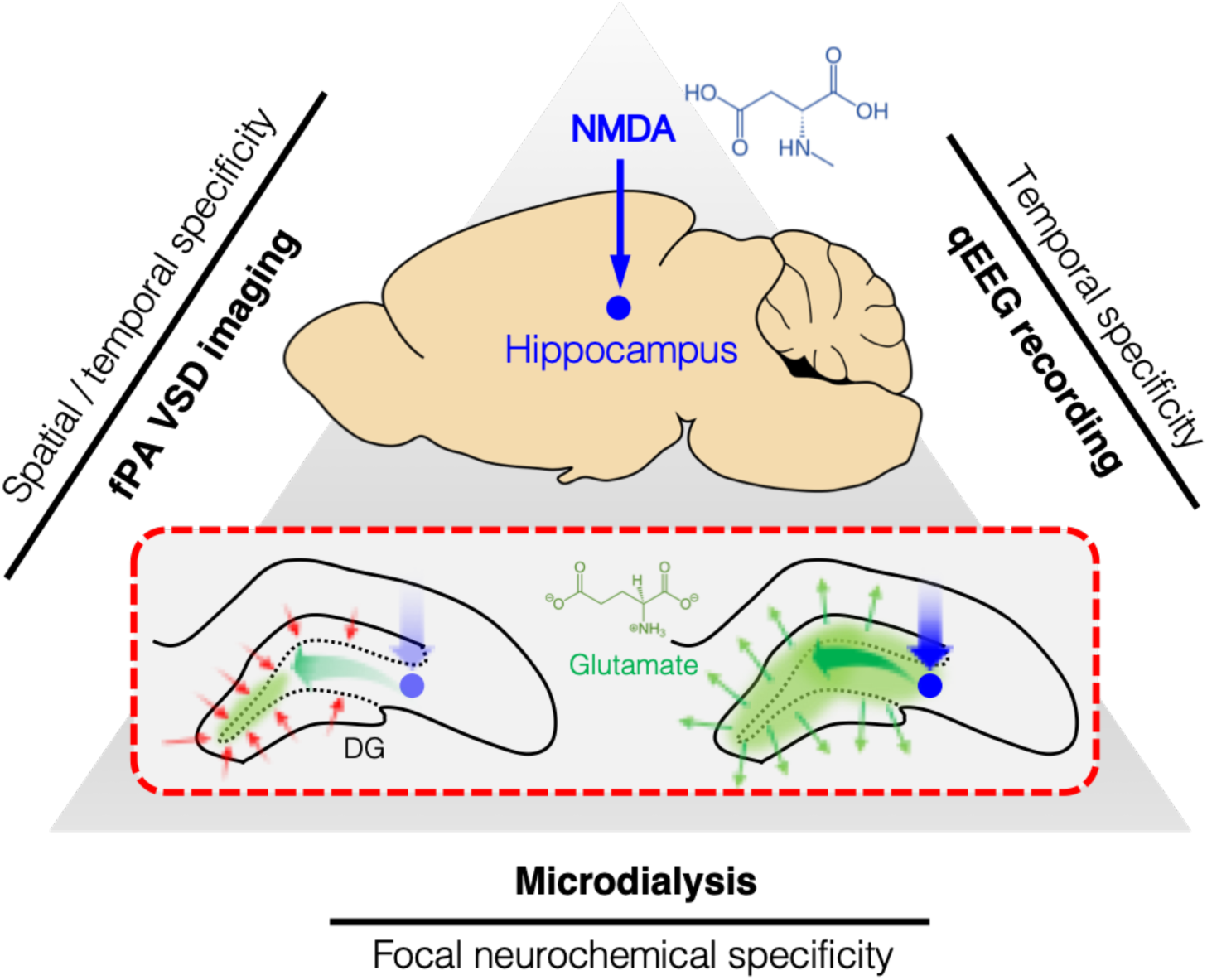
Tri-modal sensing of rat hippocampus. Red dotted rectangular describes dentate gyrus (DG) gating breakdown at hippocampus to a focal NMDA infusion. NMDA, N-methyl-d-aspartate; VSD, voltage-sensitive dye.

## Results

### Transcranial fPA neuroimaging of rat hippocampus

Transcranial fPA neuroimaging of hippocampal circuit dynamics was performed during NMDA infusion at rat hippocampus (Figure 2). The representative fPA sagittal planes obtained at 790 nm presented a sufficient sensitivity at the depths of interest, i.e., 3.6 mm, at the contralateral position to the microdialysis probe (Figure 3a). The sagittal PA imaging plane of hippocampus revealed the cross-sections of transverse sinus and inferior cerebral vein in the superficial depth range (Figure 3b). From the temporal dimension, time-averaged VSD responses were reconstructed pixel-by-pixel for pre-injection phase (−10 – 0 min) followed by the results for time bins in 10 min interval: 0 – 10 min, 10 – 20 min, and 20 – 30 min. Note that the total recording duration (40 min) was limited by internal memory of the fPA neuroimaging system. Figure 3c shows the representative VSD responses in hippocampal cross-sections collected during 0.3 mM and 3.0 mM NMDA infusion. As a result, the hippocampus with 0.3 mM NMDA infusion did not present such a significant change (*n* = 3), which implies a failure to overcome the high activation threshold of DG gatekeeping with 0.3 mM NMDA infusion. Otherwise, the substantial amount of circuit dynamics was detected at hippocampus with 3.0 mM NMDA infusion (*n* = 6). In the 3.0 mM NMDA infusion group, the peak VSD responses were presented during the first or second 10 min durations, and their peak intensities corresponds to the maximal glutamate concentration change measured in the microdialysis, e.g., 734.48 % and 493.91 % (center and right columns in Figure 3c). Further quantitative multi-modal correlation will be given in the following subsections.

**Figure 2.**
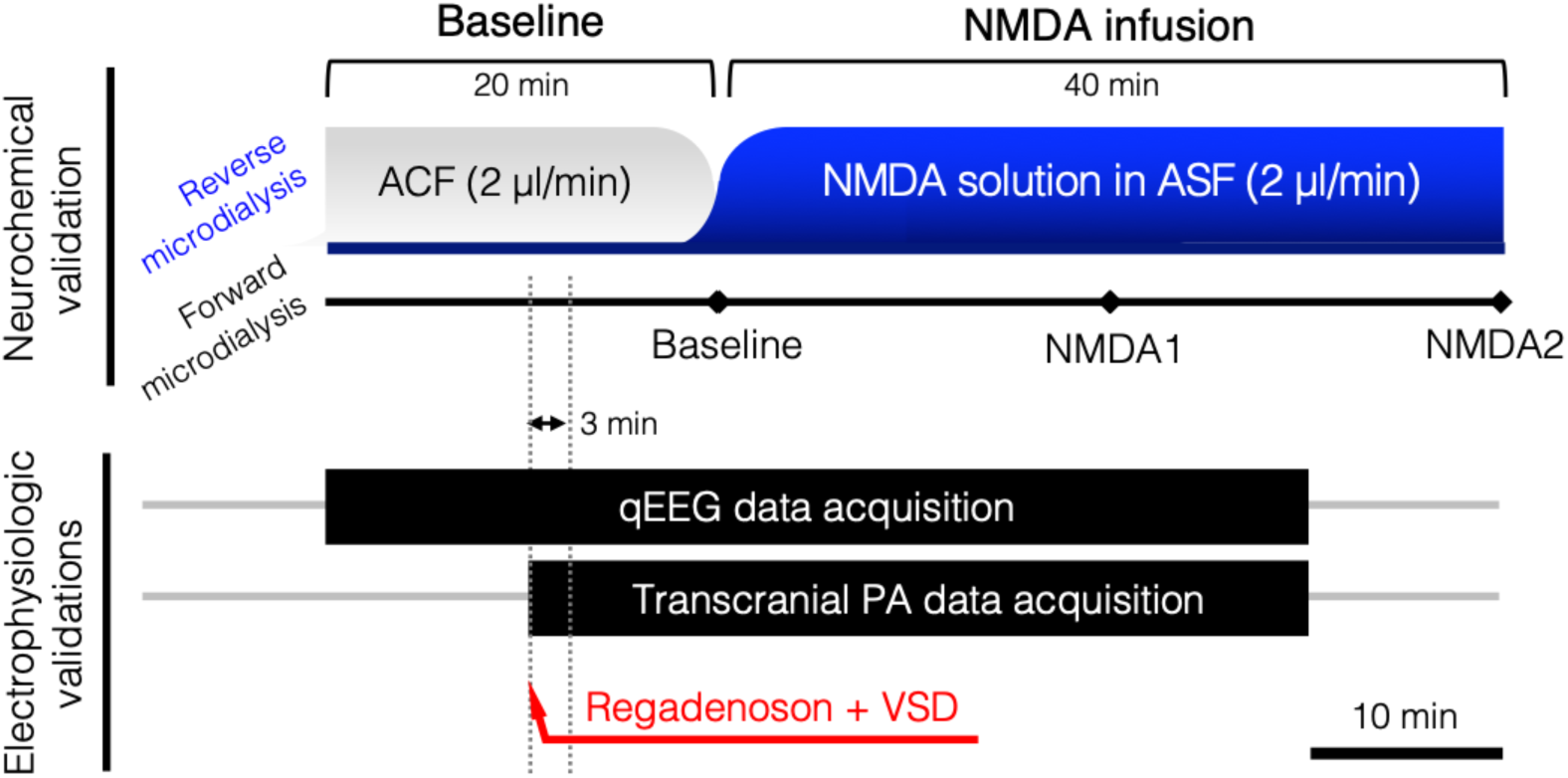
*In vivo* experimental protocol. Tri-modal monitoring of rat hippocampus using reverse/forward microdialysis, transcranial PA imaging, and qEEG. ACF, artificial cerebrospinal fluid.

**Figure 3.**
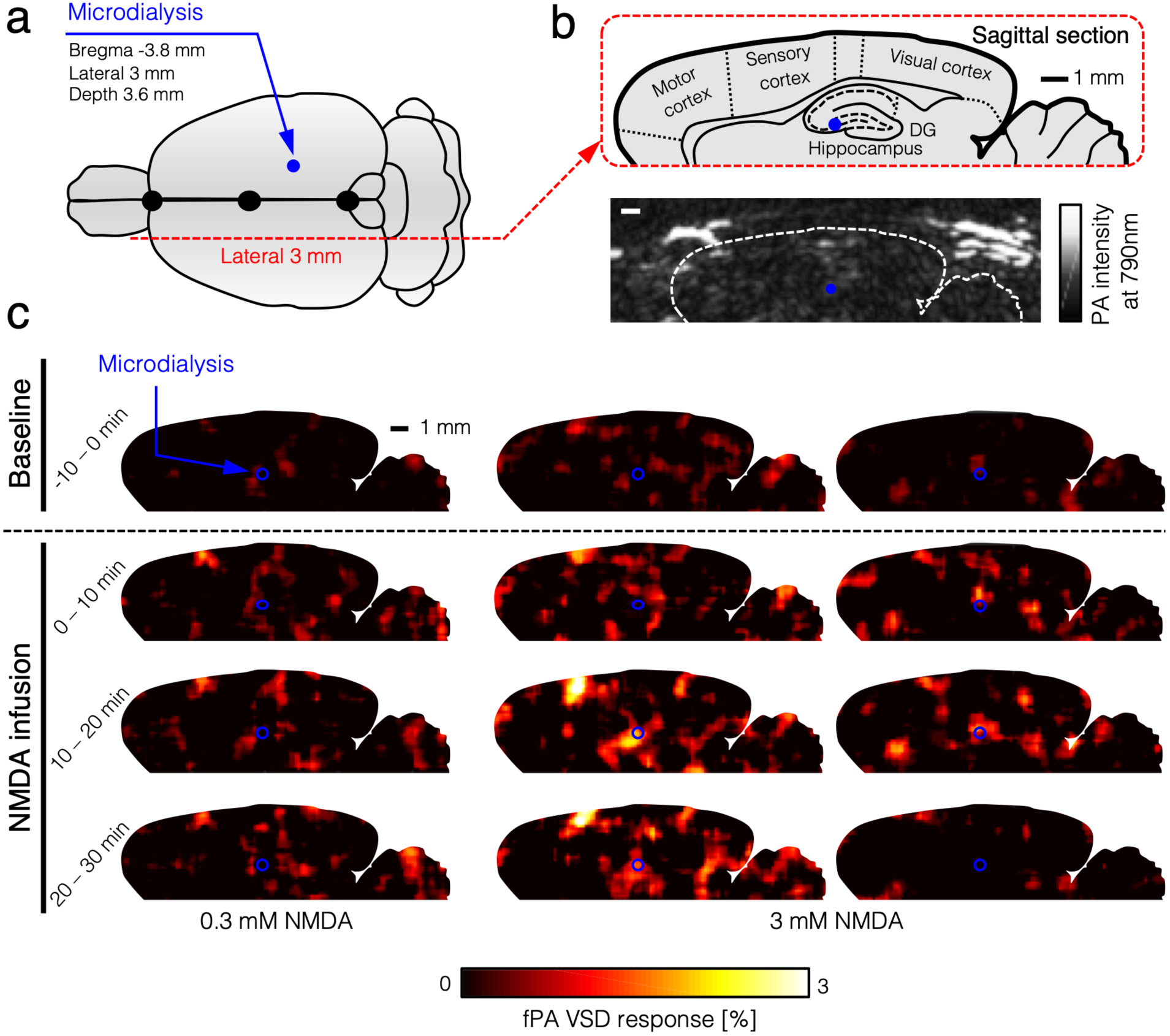
Transcranial PA neuroimaging of the hippocampal circuit dynamics following focal NMDA infusion. (**a**) Illustration of stereotaxic coordinates for the microdialysis probe and sagittal cross-sections for real-time PA recording. (**b**) The sagittal PA imaging plane was selected in the contralateral side of the microdialysis probe infusing NMDA into the brain (3 mm lateral). (**c**) Time-averaged VSD response maps during −10 – 0 min (baseline phase); 0 – 10 min, 10 – 20 min, and 20-30 min (NMDA infusion phases). Note that the blue points indicate microdialysis probe in contra-lateral positions. Maximal glutamate concentration increases for representative fPA images were 34.23 % (left), 734.48 % (center) and 493.91 % (right).

### NMDA-evoked glutamatergic neurotransmission

Intrahippocampal infusion of NMDA diluted into artificial cerebrospinal fluid at the required concentrations was performed through the microdialysis probe. Concentrations of glutamate release in the dialysate were calculated from values of three baseline samples collected at 20 min intervals before NMDA infusion. During infusion into hippocampus (bregma −3.8 mm, lateral 3 mm, depth 3.6 mm; Figure 4a), samples continued to be collected every 20 min for the remainder of the study. Glutamate release was calculated as a % of basal values. Figure 4b shows the effect of NMDA infusion on glutamate release in the hippocampus of anesthetized rats. The results demonstrate the dose-related response of NMDA infusion at 0.3 (*n* = 3), 1.0 (*n* = 1), and 3.0 mM (*n* = 6) into the hippocampus, where 0.3 mM NMDA caused a 17.93±19.05 % increase in glutamate levels that remained elevated for the duration of the infusion; the 1.0-mM NMDA infusion appears to cause doubling of the glutamate levels in the hippocampus compared to baseline (i.e., 112.42 %). This increase observed at the 1.0-mM remained elevated through to the end of the NMDA infusion. However, at 3.0-mM, the NMDA infusion raised the glutamate level up to 392.99±250.54 %, above baseline level, which peaked during the second 20-min sampling period.

**Figure 4.**
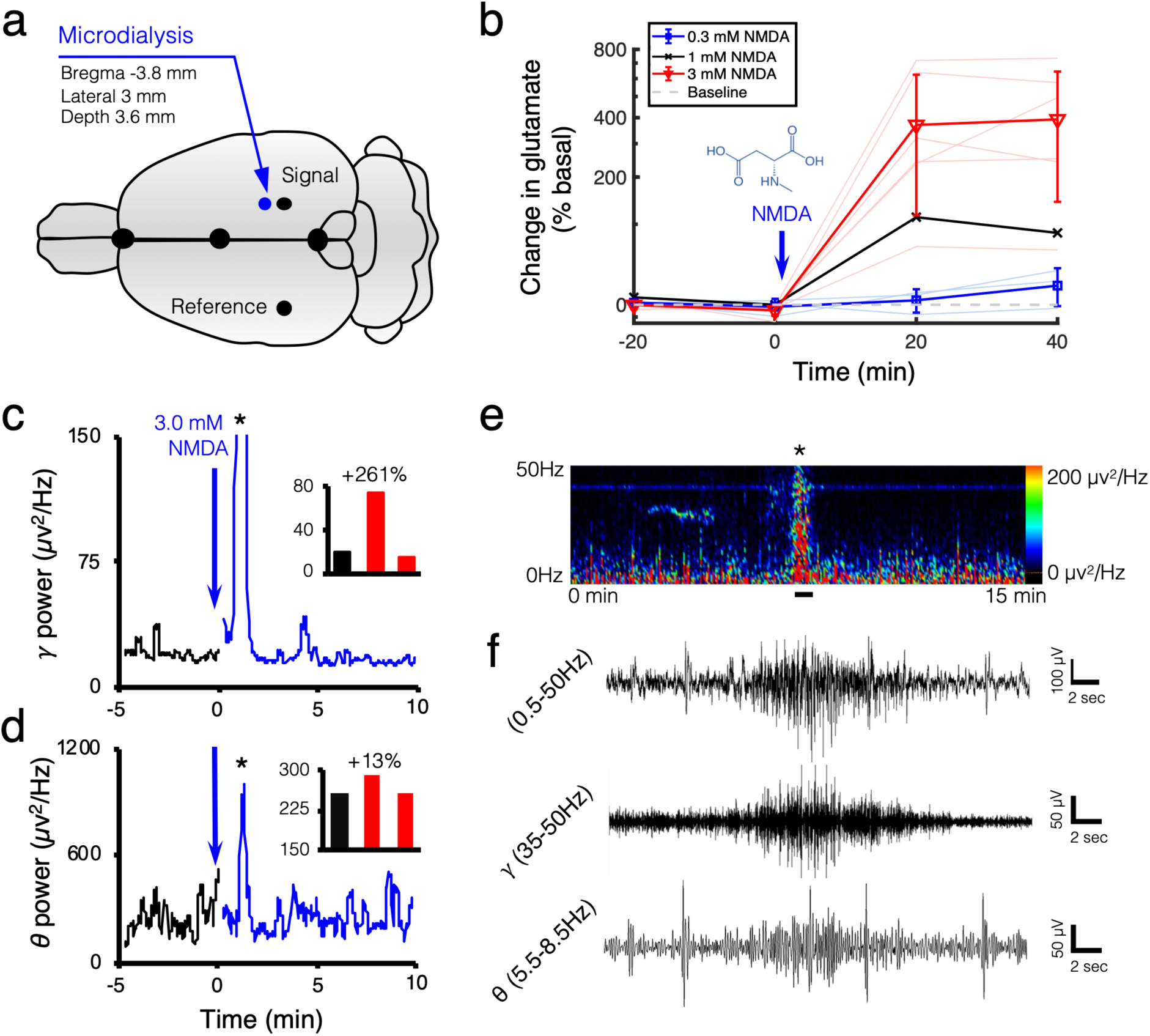
Extracellular glutamate concentration and concomitant electrophysiology during NMDA-evoked activity in the hippocampus. (**a**) Illustrates stereotaxic coordinates for the microdialysis probe and recording leads. Ground electrode was implanted over the rostrum. (**b**) 0.3 (*n* = 3), 1.0 (*n* = 1), and 3.0 mM (*n* = 6) NMDA infusion into the hippocampus caused maximal glutamate increases of 17.93±19.05%, 112.42%, and 392.99±250.54% respectively as compared to % baseline. Gray dotted line indicates the baseline. (**c**) Gamma (35-50Hz) power shows a 5-min baseline recording (black trace), followed by two consecutive 5 min traces (blue trace) following focal NMDA infusion in the hippocampus. EEG demonstrated a 261% increase in gamma power, as compared to % baseline, and an onset of epileptiform discharges after focal NMDA infusion (see inset bar graph; asterisk denotes epileptiform activity). (**d**) Respective theta power after NMDA infusion represents the theta component of the focal seizure. (**e**) 15-min spectral power heat map demonstrates the spectral power changes associated with NMDA infusion and subsequent epileptiform activity (denoted by black asterisk in e). (**f**) The representative raw EEG trace during the occurrence of the epileptiform event (solid black line in e) for full (05.-50Hz), gamma (0.5-50Hz), and theta (5.5-8.5Hz) power; respectively. For the expanded time scale of the focal hippocampal seizure see Supplemental 1.

### EEG of electrophysiological activity in hippocampus

The DG is a hippocampal region specifically subjected to a barrage of excitatory inputs. However, the majority of excitatory activity does not propagate through the DG and into the hippocampus as the DG performs a gating function ^1^. Excessive activation of the DG disrupts its gating function and induces acute seizures in naïve animals ^1^. Hippocampal infusion of 3.0 mM NMDA induced a maximal VSD and significant rise in extracellular glutamate concentration, suggesting a potential break in the DG gate that may lead to acute seizure activity. We therefore utilized qEEG during the same hippocampal 3.0 mM NMDA infusion protocol to identify if the maximal VSD response and extracellular glutamate concentrations were associated with significant electrophysiological activity, a tri-modal approach that validates the temporal and spatial resolution of fPA imaging during focal NMDA infusion.

In patients with focal epilepsy, gamma and theta activity from scalp EEG are an indicator of the seizure onset zone and ictal onset ^20,21^. Here, the temporal specificity of qEEG recorded circuit responses in real-time, before and after focal NMDA infusion. The 3.0 mM NMDA infusion into the hippocampal circuit induced focal seizure activity recorded on qEEG (Figure 4c black asterisk and S1). When NMDA was delivered to the hippocampal circuit by the microdialysis probe (Figure 4a)^22^, a significant change in qEEG spectral power was presented, especially in the gamma range (Figure 4c). In the theta range, spectral power increased during the seizure (Figure 4d) further supporting the identification of the focal seizure activity. The seizure trace on EEG demonstrated both a gamma and theta component (Figure 4f). Importantly, the ictal event occurred during the same temporal window as the maximum VSD response and the greatest increase in extracellular glutamate concentration. The dose-related VSD and extracellular glutamate concentration response with qEEG suggest that the 3.0-mM NMDA infusion resulted in a break in the DG gate associated with focal seizure activity.

However, qEEG lacks spatial resolution, and recordings from deep brain structures cannot be isolated without implantation of invasive depth electrodes. Therefore, in order to maintain the tri-modal experimental paradigm we opted not to place depth electrodes. We made the stimulation focal, according to the previous proof-of-concept results by fPA detection onset of generalized seizures through an intact skull and scalp ^19^.

### Quantitative multi-modal correlation

The fPA VSD responses at the contralateral side to the microdialysis probe (blue dots in Figure 3c) were measured and plotted as a function of the corresponding changes in extracellular glutamate concentration (Figure 5). Note that the regions-of-interest ROIs to quantify the VSD responses were in 1 x 1 mm^2^ size in the sagittal cross-section of rat hippocampus. The reference phase for VSD response quantification was obtained from the baseline phase: 5 – 10 min. In fPA imaging, 3.0 mM NMDA infusion at hippocampus yielded significant elevation of VSD response: 0.08±0.35, 2.24±1.06, and 1.17±1.56 for baseline (−10 – 0 min), NMDA1 (0 – 20 min), and NMDA2 (20 – 30 min) phases, respectively (Figure S2a). The increase of glutamate concentration change was correspondingly presented from −4.67±4.48 %, 369.75±254.91 %, to 392.99±250.54 %. In addition, when re-analyzing the data for −50 – 100 %, 100 – 500 %, and 500 – 1,000 % bins of fractional glutamate concentration change, high positive correlation was obtained with VSD responses: 0.38±0.55 (*n*_sample_ = 9), 0.97±1.40 (*n*_sample_ = 5), and 3.17±0.32 (*n*_sample_ = 4) for 11.04±29.60 %, 279.85±141.64 %, to 666.49±73.63 % of glutamate concentration changes (red lines in Figure 5). The goodness of fit (R^2^) among the mean values was 0.95 with slope and y-intercept at 0.12 and 0.00, respectively. The qEEG found the increased bursts of gamma power during 3.0 mM NMDA infusion, reaching up to ∼261 % increase with a focal seizure (Figure 4c and S1). Otherwise, focal 0.3 mM NMDA infusion at hippocampus did not presented any circuit activity increase statistically significant. The VSD responses were −0.11±0.02, −0.15±1.71, and - 0.07±0.64 in baseline, NMDA1, and NMDA2 phases, respectively, which are correlated to the low glutamate concentration changes in hippocampus: −1.65±6.90 %, 3.90±10.41 %, and 17.93±19.05 % (Figure S2b). Also, the VSD responses presented strong positive correlation when re-analyzed in −10 – 0 %, 0 – 10 %, and 10 – 50 % bins of fractional glutamate concentration changes: −1.00±1.00, 0.10±0.35, and 0.58±0.48 (*n*_sample_ = 3 for each) with −6.84±3.38 %, 4.43±4.05 %, and 22.59±11.64 % of fractional glutamate concentration changes, respectively (blue lines in Figure 5). The R^2^ was 0.88 with slope and y-intercept at 0.05 and −0.46, respectively.

**Figure 5.**
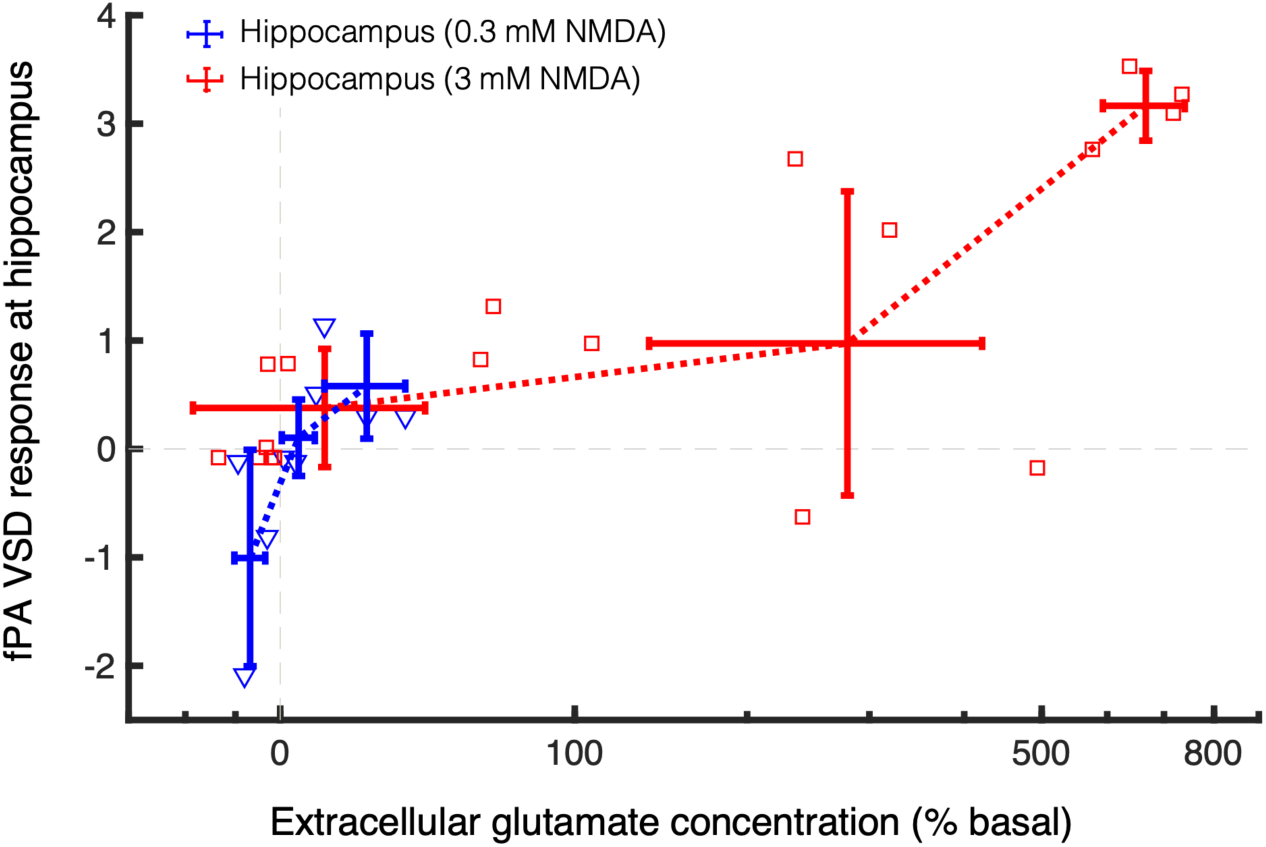
fPA VSD response at hippocampus as a function of extracellular glutamate concentration change. Grey dotted lines indicate the basal level in fPA VSD response and extracellular glutamate concentration.

### Brain histology

Brain tissue was extracted after all *in vivo* experimental protocols. The brain was frozen-sectioned into 300-µm thickness slices and evaluated to confirm probe placement using the hemorrhage caused by the microdialysis probe insertion for the NMDA infusion and collections as a marker. Figure 6a shows the bright-field images of the coronal plane of the hippocampus at −3.8 mm from bregma, respectively. White arrows indicate the hemorrhage caused by the microdialysis probe positioned at the following coordinates in hippocampus: bregma −3.8 mm, lateral 3 mm, and depth 3.6 mm. The VSD staining of brain tissue was confirmed with near-infrared fluorescence microscopy. Uniform VSD fluorescence was found in the VSD perfusion animal, while negative control (VSD-) presented negligible fluorescence emission (Figure 6b), which again confirms the results in our previous publication ^23^.

**Figure 6.**
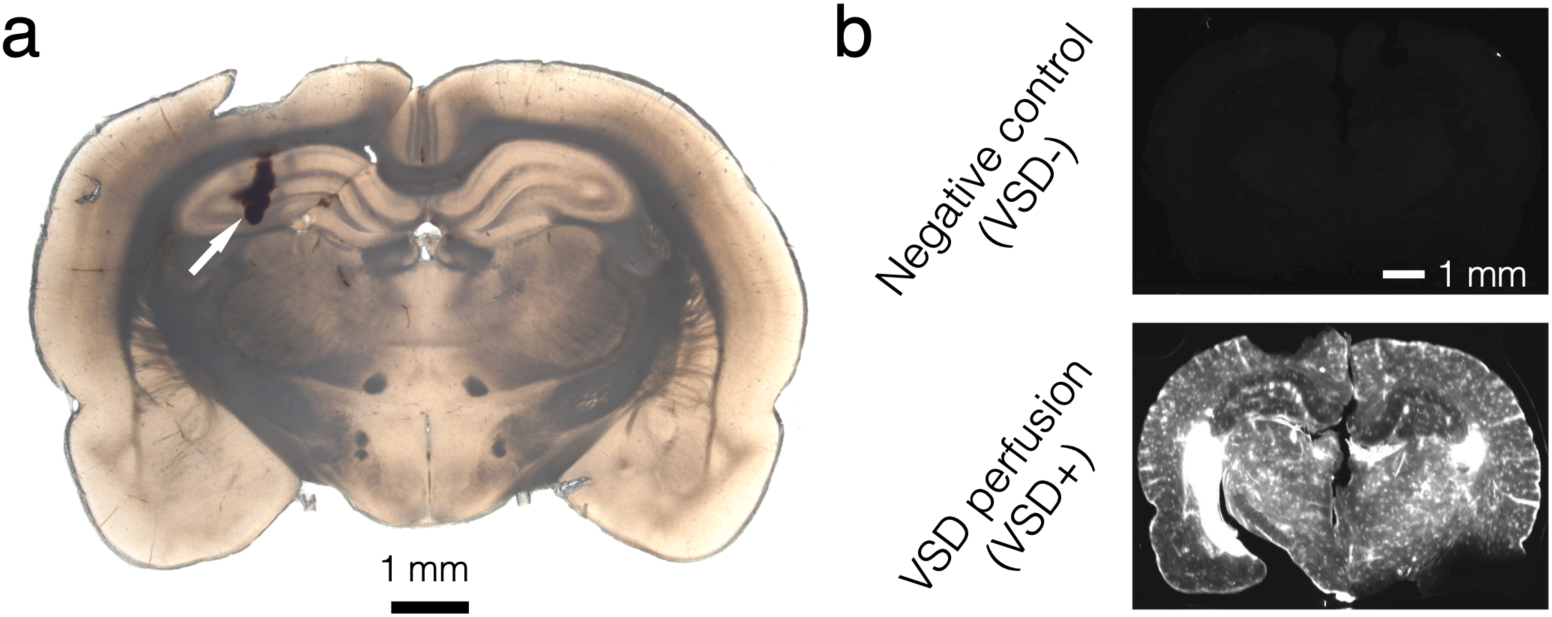
Histopathological confirmation. (**a**) Microdialysis probe at hippocampus. White arrow indicates the wound caused by microdialysis probe installation. (**b**) Frozen-sectioning histopathological confirmation of systematic VSD delivery throughout brain tissue region.

## Discussion

The application of fPA neuroimaging was expanded to neuroscience ^24-26^ as label-free transcranial fPA imaging of neurovascular coupling proposed as a means to quantify hemodynamic changes ^27,28^. However, this approach does not yield quantitative neural activities that directly correspond to electrical activity. Hemoglobin provides an effective contrast signal in fPA neuroimaging, but the neurovascular coupling in brain is comparatively slow compared to electrophysiological neural activities. Instead, there has been extensive investigations into more effective exogenous contrast agents ^29-33^. This approach has enabled several neuroimaging approaches with functional voltage sensors. Deán-Ben et al. showed real-time PA tomography of a genetically encoded calcium indicator, GCaMP5G, using zebrafish *in vivo* ^34^. Sheryl Roberts, et al. also proposed a new metallochromic calcium sensor for PA imaging (CaSPA) which has a high extinction coefficient, a low quantum yield, and high photo-bleaching resistance for brain and heart imaging ^35^. Ruo et al. reported PA imaging of neural activity evoked by electrical stimulation and 4-aminopyridine-induced epileptic seizures using hydrophobic anions such as dipicrylamine (DPA) in mouse brain ^36^. However, these voltage sensors requires PA imaging at the visible spectral range (488 nm and 530 nm for GCaMP5G; 550 nm for CaSPA; 500 nm and 570 nm for DPA), which are suboptimal when imaging deep brain such as hippocampus positioned at 5 mm – 8 mm depth including intact scalp and cortex in rat ^22,37^.

Recently, we proposed transcranial fPA recordings of brain activity *in vivo* with a near-infrared VSD, delivered through the blood-brain barrier (BBB) via pharmacological modulation, as a promising tool to transcend optical neuroimaging limitations, particularly as it relates to sensing depth ^19,23,38^. The studies demonstrated that transcranial fPA neuroimaging distinguishes *in vivo* seizure activity in stimulated rat brains from that of control groups in real time. However, the results were limited by the use of a global chemo-convulsant, causing perturbation across the entire brain caused by the intraperitoneal administration of penetylenetetrazole (PTZ). In this paper, we presented follow-on advances in fPA VSD neuroimaging by focal neural stimulation of heterogeneous neural circuits, with concomitant validations from qEEG and glutamate quantification using microdialysis, respectively. The set of experiments described here yield key findings as follows: (1) The microdialysis-dependent low- and high-dose NMDA infusion into the central nervous system (CNS) lead to a wide range of focal extracellular glutamate concentration increase in hippocampus up to ∼800 %. (2) The neurochemical response (microdialysis) was well-correlated to the phenotypes in the electrophysiological sensing (qEEG). The NMDA activation in the hippocampus triggers an all-or-none type of circuit dynamics that lead to the initiation of a focal seizure in the hippocampal circuit. (3) Transcranial fPA neuroimaging data successfully identified hot spots of focal NMDA receptor activation, as presented in the qEEG recordings. The hippocampal circuitry provided the proportional excitation of glutamatergic neurotransmission with concomitant NMDA infusion. The DG of the hippocampus is positioned as a gatekeeper to regulate the vast excitatory cortical inputs from propagating into the hippocampus ^1^. Characteristically, the DG displays a high activation threshold; a trait that is mediated by its profuse innervation by inhibitory GABAergic neurons and relatively hyperpolarized resting membrane potential of its pyramidal neurons ^39^. In epilepsy the DG fails to gate the propagation of excitatory inputs into the hippocampus, resulting in overexcitation and seizures. In naïve animals, *in vivo* optogenetic activation of the DG disrupts its gating function and induces seizures that increase in severity depending on the duration of the stimulus ^1^. In this study, the disruption of DG gating by strong stimuli has been clearly demonstrated by utilizing the fPA VSD neuroimaging techniques during focal NMDA infusion.

Further investigations are required to advance our current perspectives available with tri-modal sensing, including fPA, qEEG, and microdialysis. Glutamate produces fast-rising brief depolarizations in pyramidal neurons. Therefore, the use of fPA neuroimaging will enable us to more precisely assess glutamate and GABA dynamics in order to formulate a more complete profile of circuit activation. Once homeostasis is disrupted, neuronal activity is sensitive to changes both of excitatory and inhibitory mechanisms. Faster neurochemical recording is another approach that may prove useful in assessing the impact of these measures. Although microdialysis successfully provided quantitative, focal neurochemical concentrations, the sampling rate of 1 sample per 20 min was slow. Techniques offering faster temporal resolution may allow more meaningful comparison of the neurochemical changes yielded by microdialysis and the electrophysiological events monitored by qEEG and fPA neuroimaging ^40^. One such technique, using custom built hardware and the Amplex Red method, achieved fluorescence-based quantification of glutamate from samples taken every 5 seconds, though reliability appeared to be limited when higher glutamate concentrations were measured ^41^. Implantable glutamate biosensors allow sub-second readouts of neurochemical concentrations. However, current limitations include sensitivity, selectivity, and high cost; recent developments in materials, effective modeling, and sensor design may soon alleviate some of these limitations ^42,43^.

From the results, transcranial fPA neuroimaging was able to differentiate the circuit activity defined with qEEG and microdialysis. However, future developments should serve to further advance the efficacy of the fPA neuroimaging in neuroscience. (1) We expect that improved signal processing for extracting neural activity from the ubiquitous blood context will enable better characterization of brain function. The present *in vivo* experiments confirmed the possibility of background suppression, as also presented in our previous study ^19^. Enhanced signal processing and/or use of multi-spectral wavelengths may allow significantly improved spectral differentiation of electrophysiological activities in the brain at higher temporal resolution, leading to development of novel quantitative metrics for real-time brain activity measures. Having isotropic resolution with 2-D PA probes would be also an interesting direction to pursue as a follow up to the present work. The use of 2-D PA probe would not only allow real-time volumetric information, but also enable the suppression of off-axis interference. Even though we presented that neural activity can be successfully discerned with our current 1-D PA probe, its sensitivity might be affected by off-axis interferences especially from the elevation direction because of the limited acoustic lens focusing at a fixed depth. The neuroimaging using 2-D PA probe would reject those interferences by the advanced electrical beamforming capability in axial, lateral, and elevation directions. Having an improved PA imaging system would provide significant breakthrough in terms of spatiotemporal resolution in fPA neuroimaging. Even though our current laser system yields both 4 fps (frame-per-second) of temporal resolution and PA signal sensitivity at rat hippocampus, further optimization of temporal resolution would provide finer spatiotemporal specificity. On the other hand, we consider employing larger animal model for this fPA VSD neuroimaging research. Pig models have been an ideal subject to pave the way to human translation of neuroengineering technologies, thanks to their analogous brain structure and physiology and scalp and skull thicknesses to those in humans, with alleviated ethical issues ^44^. We already validated the transcranial fPA neuroimaging in the pig model, and will continue to pursue this research direction ^45^.

In all, the transcranial fPA neuroimaging at hippocampus in *in vivo* rat brain was successfully correlated with electrophysiologic and neurochemical measurements using qEEG and microdialysis: focal NMDA infusion triggers glutamate release that excites the neural circuit, and at threshold doses it causes runaway excitation in the hippocampus by overcoming DG gating. This is reflected in the lower seizure threshold of the hippocampus. Therefore, the transcranial fPA neuroimaging is a promising technology for the visualization of focal neural events in real time.

## Material and Methods

### Animal preparation

For the proposed *in vivo* experiments, 8-9-week-old male Sprague Dawley rats (Charles Rivers Laboratory, Inc., MA, United States) weighing 275-390g were used. The use of animals for the proposed experimental protocol was approved by the Institutional Animal Care and Use Committee of Johns Hopkins Medical Institute (RA16M225). Rats were housed in groups of 3 per cage with free access to food and water and maintained on a 12hr light / 12hr dark cycle.

On the day of the study the rats were weighed and anesthetized with urethane (1.2mg/kg,). Urethane was given incrementally with alternating intra-peritoneal (ip) and subcutaneous (sc) dosing. Three (3) ml of isotonic saline was given sc on each side of the body to keep the animal hydrated during the experimental procedure. Body temperature was maintained until animal was fully anesthetized and ready for surgery. For fPA and qEEG studies, an iv catheter was inserted into a tail vein prior to surgery for dye administration during the studies. Once a stable plane of anesthesia was established, hair was shaved from the scalp of each rat to have acoustic coupling for transcranial fPA recording. The rat was placed into a stereotaxic device (Stoeling Co. Wood Dale, IL). This fixation procedure was required to prevent any unpredictable movement during fPA or EEG recording of neural activities. A CMA12 microdialysis probe (Harvard Apparatus, Holliston, MA, USA) was implanted into the CA_3_ region of the right hippocampus (stereotaxic coordinates: 3 mm lateral and 3.8 mm posterior to bregma, and 3.6 mm below the surface of the dura, Figure 3a) ^22^. The probe active exchange surface was 2 × 0.5 mm. The probe was secured to the skull using dental acrylic cement. The fPA and qEEG probes were placed on the contralateral side of the microdialysis probe.

### Fluorescence quenching-based near-infrared voltage-sensitive dye

In the present *in vivo* study, we used the fluorescence quenching-based near-infrared cyanine VSD, IR780 perchlorate (576409, Sigma-Aldrich Co. LLC, MO, United States) as used in our previous *in vivo* study differentiating a chemo-convulsant seizure activity ^19^, and it has the analogous chemical structure of PAVSD800-2, our new VSD validated in our previous *in vitro* study ^38^. This VSD yields fluorescence quenching and de-quenching depending on membrane polarization and subsequent change in the local VSD molecule density, leading to a reciprocal change of PA contrast with non-radiative relaxation of absorbed energy.

### Functional fPA neuroimaging

We used real-time PA data acquisition to record electrophysiological neural activities *in vivo* as in our previous study ^19^: an ultrasound research system consisted of an ultrasound linear array transducer connected to a real-time data acquisition system (SonixDAQ, Ultrasonix Medical Corp., Canada). To induce the PA signals, pulsed laser light generated by a second-harmonic (532 nm) Nd:YAG laser pumping an optical parametric oscillator (OPO) system (Phocus Inline, Opotek Inc., USA) provided 690-900 nm of tunable wavelength range and 20 Hz of the maximum pulse repetition frequency. A bifurcated fiber optic bundle, each 40 mm long and 0.88 mm wide, was used for laser pulse delivery. The PA probe was situated between the outlets of the bifurcated fiber optic bundles using a customized, 3-D printed shell for evenly distributing laser energy density in the imaging field-of-view. The alignment of outlets was focused specifically at 20 mm depth. The PA probe was positioned in the contralateral sagittal plane of microdialysis probe (3 mm) to cover the hippocampal cross-section. The distance between the PA probe and the rat skin surface was 20 mm filled with acoustic gel, and the resultant energy density was at ∼3.5 mJ/cm^2^, which is far below the maximum permissible exposure (MPE) of skin to laser radiation by the ANSI safety standards ^46^. A wavelength of 790 nm was used, at which sufficient absorbance can be obtained by the near-infrared VSD, i.e., IR780 perchlorate. Also, excitation at that wavelength prevented the undesired time-variant change of blood oxygen saturation, since the wavelength corresponds to the isosbestic point of Hb and HbO_2_ absorption spectra. Detailed information of neural activity reconstruction using normalized time-frequency analysis can be found in our previous publication ^19^.

### *In vivo* microdialysis

*In vivo* microdialysis sampling was carried out as previously described ^47,48^. For infusion experiments, NMDA (Sigma-Aldrich Chemicals, St. Louis, Mo) was weighed, solubilized, and diluted to the desired concentration in artificial cerebrospinal fluid (NaCl, 147 mmol/L; KCl, 2.7 mmol/L; CaCl_2_, 1.2 mmol/L; MgCl_2_, 0.85 mmol/L) (Harvard Apparatus, Holliston, MA, USA) on the study day. Once the probe was inserted and secured, it was perfused with artificial cerebrospinal fluid pumped at a flow rate of 2 μl/min. Samples were collected at 20 min intervals, and immediately transferred to a −80°C freezer until assayed. To allow sufficient time for the glutamate levels to equilibrate, three baseline samples were collected an hour following initiation of infusion. Following these samples, NMDA was infused into the brain directly through the dialysis probe with the same pump parameters as used for the baseline samples. Dialysate samples were assayed for glutamate by a two-step process using HPLC-ECD on an Eicom HTEC-500 system (EICOM, San Diego, CA, USA). After passing the samples through a separation column, they were processed via a column containing immobilized L-glutamate oxidase enzyme, resulting in the release of hydrogen peroxide. The hydrogen peroxide concentration was then determined using a platinum working electrode. Chromatographic data were acquired online and exported to an Envision software system (EICOM, San Diego, CA, USA) for peak amplification, integration, and analysis.

### Quantitative EEG

All EEG recordings utilized a three-electrode paradigm: 1 recording, 1 reference (aligned to the site of activation) and 1 ground over the rostrum. The electrodes (IVES EEG; Model # SWE-L25 – IVES EEG solutions, MA, USA) were fixed with minimal cyanoacrylate adhesive (KrazyGlue), similar to previous protocols ^49^. Data acquisition was performed using Sirenia software (Pinnacle Technologies Inc., Kansas, USA) with synchronous video capture. Data acquisition had a 14-bit resolution, 400 Hz sampling rate, and a band pass filter between 0.5 Hz and 50 Hz. The acquisition files were stored in an .EDF format and scored manually, using real-time annotations from the experiments. EEG power for 2-second epochs was done using an automated fast Fourier transformation module in Sirenia software ^50^.

### *In vivo* experimental protocol

The *in vivo* protocols were designed for simultaneous multi-modal sensing of the neural activity at hippocampus: microdialysis-qEEG and microdialysis-fPA neuroimaging. Figure 2 shows a detailed schematic protocol for each group representing the response to the administration of NMDA, Lexiscan and VSD (i.e., IR780 perchlorate). fPA and qEEG data acquisition were performed for 40 min to correlate with three microdialysis samples collected at 20-min intervals. Graded NMDA infusion concentrations were applied to identify the dose-dependent glutamatergic excitation of hippocampal circuit: 0.3 mM (*n* = 3) and 3.0 mM (*n* = 6). VSD and Lexiscan followed the data acquisition sequence with 3-min delay, thereby 5 min of baseline phase was guaranteed for the VSD response reconstruction in fPA neuroimaging before starting NMDA infusion. The dosing protocol for Lexiscan and VSD administration was as follows: through an iv tail vein catheter, 150 µl of Lexiscan (0.4mg/5ml) was injected, followed by 200 µl of VSD at 2 mg/ml concentration, flushed immediately with 150 µl of 0.9% isotonic saline. The EEG signal was recorded an identical preparation procedure as the fPA neuroimaging, including animal preparation and administration of IR780, Lexiscan and experimental duration time for all recordings.

### Brain histology

Rats used for the above protocol were sacrificed, and whole brains immediately harvested and placed in 10 % formalin. All brains were allowed to fix in fresh 10 % formalin for at least 48 hours with gentle agitation on a conical rotator. Subsequently, the brains were processed through a series of sucrose gradients (15 %, 20 %, 30 % for 12-24 hours each) for cryoprotection. Brains were sectioned frozen at 300 µm thickness. Tissue sections were mounted on slides in ProLong Diamond Anti-face mountant. Slides with sections were imaged using an Olympus OM-D E-M5 Mark II for bright field image and using LI-COR Odyssey for fluorescence visualization.

## Author Contributions

DFW, AAG and MB originally conceived of the NMDA administration and of microdialysis with PA idea and helped to design the overall research plan with initial funding for the *in vivo* experiments. DFW critically revised the focus of the results and final version and interpretation. EMB, LML, and SDK helped to design the overall research plan, helped plan specific experiments and all contributed to the review and writing of the manuscript. Jeeun K, SDK, JSE, BJS, HV planned and carried out *in vivo* experiments, analyzed the research outcomes, and wrote key elements of the first draft of the manuscript. Jeeun K analyzed and interpreted the PA measurements and completed the first manuscript, HV and JE the NMDA dosing and microdialysis, and SDK and BS the EEG experiments. MB provided resources and personnel and vital collaboration for the microdialysis part of the experiment. HV and LML devised the VSD vehicle preparation. APM performed histopathological analysis. MMH planned and supervised confirmation of VSD penetration into brain tissue also contributed to the final version of the manuscript. AAG critically revised experimental design, draft, and final versions of manuscript, and interpretation of results. Jin K developed, funded, and participated in the current PA system design. AR participated in early planning and critically read and edited the manuscript. He has contributed both technically and materially to support this research. EMB led the development, system specification, design specification, and funding of the current PA imaging system. Secured the funding of the needed imaging experiments throughout the lifetime of the project, including taking responsibility of 2 full-time research members specifically for this work, Jeeun K and APM. Further, he has contributed intellectually by mentoring these members and providing input on manuscripts, and participating on PI meetings.

## Supplementary Material

**Figure S1.**
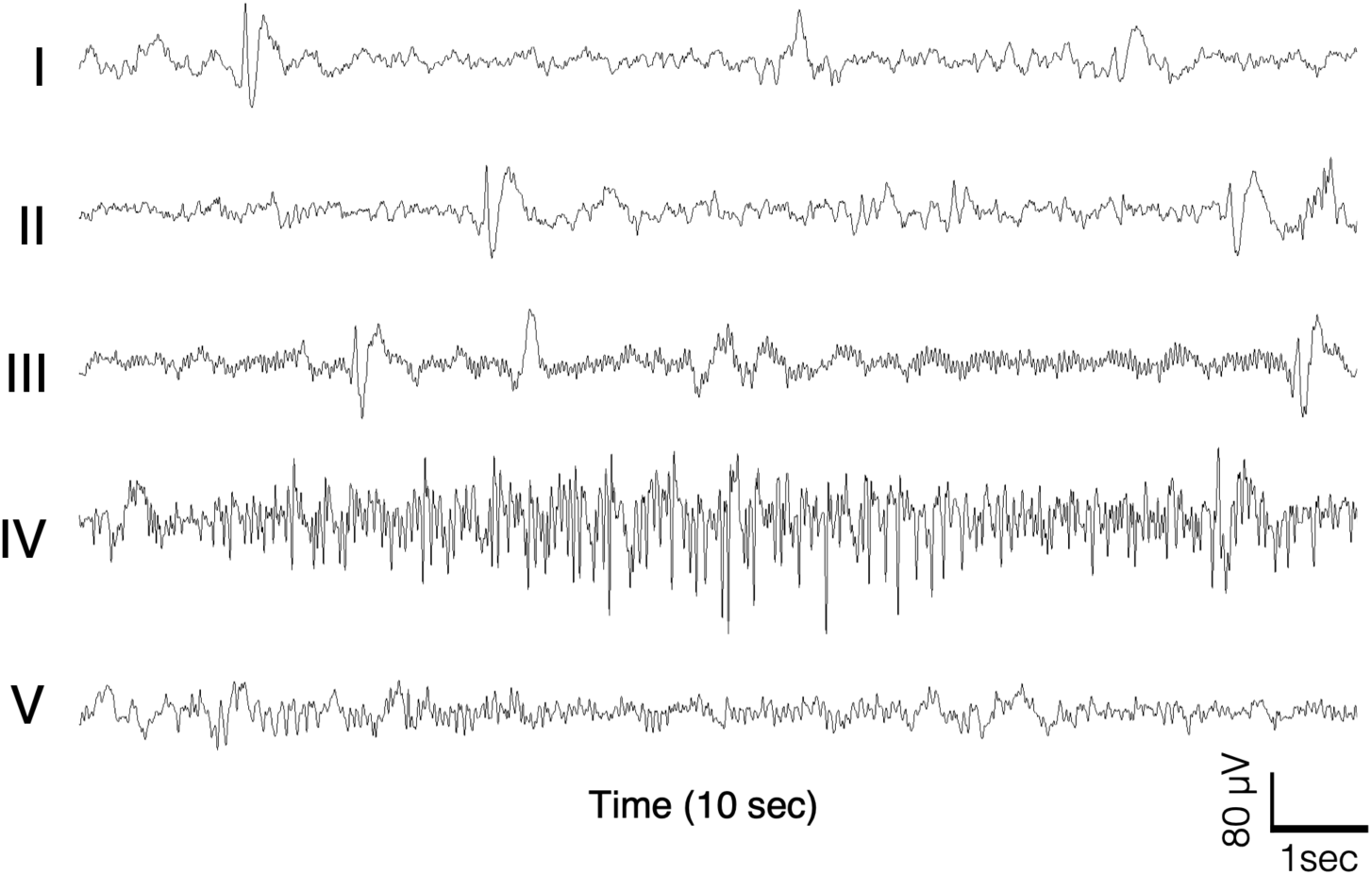
Expanded time scale of EEG recording during NMDA infusion into the hippocampus. (**a**) 10 sec expanded time scale of raw EEG traces during hippocampal NMDA infusion. (I) Baseline (II) immediately after onset of NMDA infusion (III) immediately before ictal event (IV) during focal seizure event (V) and sustained short duration high frequency hippocampal discharges after the focal seizure event.

**Figure S2.**
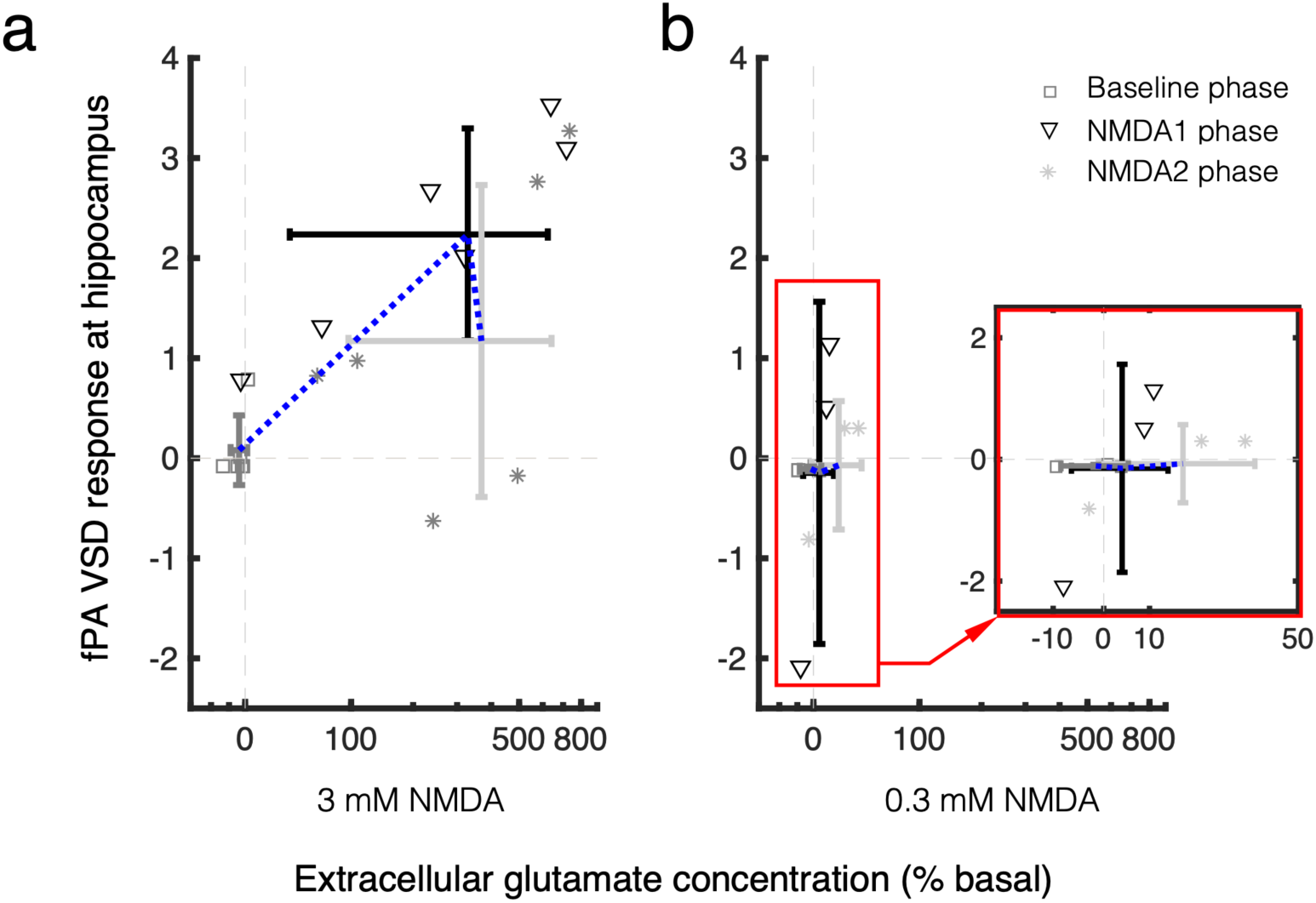
fPA VSD responses in baseline, NMDA1, and NMDA2 phases as a function of extracellular glutamate concentration change at hippocampus. (**a**) 3.0 mM NMDA infusion. (**b**) 0.3 mM NMDA infusion. 0.3 mM NMDA data in red rectangular is magnified and presented together (see inset graph). Grey dotted lines indicate the basal level in fPA VSD response and extracellular glutamate concentration.

